# Rare Genetic Variants Underlie Outlying levels of DNA Methylation and Gene-Expression

**DOI:** 10.1101/2020.02.19.950659

**Authors:** V. Kartik Chundru, Riccardo E. Marioni, James G. D. Pendergast, Tian Lin, Allan J. Beveridge, Nicholas G. Martin, Grant W. Montgomery, David A. Hume, Ian J. Deary, Peter M. Visscher, Naomi R. Wray, Allan F. McRae

## Abstract

Testing the effect of rare variants on phenotypic variation is difficult due to the need for extremely large cohorts to identify associated variants given expected effect sizes. An alternative approach is to investigate the effect of rare genetic variants on low-level genomic traits, such as gene expression or DNA methylation (DNAm), as effect sizes are expected to be larger for low-level compared to higher-order complex traits. Here, we investigate DNAm in healthy ageing populations - the Lothian Birth cohorts of 1921 and 1936 and identify both transient and stable outlying DNAm levels across the genome. We find an enrichment of rare genetic variants within 1kb of DNAm sites in individuals with stable outlying DNAm, implying genetic control of this extreme variation. Using a family-based cohort, the Brisbane Systems Genetics Study, we observed increased sharing of DNAm outliers among more closely related individuals, consistent with these outliers being driven by rare genetic variation. We demonstrated that outlying DNAm levels have a functional consequence on gene expression levels, with extreme levels of DNAm being associated with gene expression levels towards the tails of the population distribution. Overall, this study demonstrates the role of rare variants in the phenotypic variation of low-level genomic traits, and the effect of extreme levels of DNAm on gene expression.

## Introduction

DNA methylation (DNAm) is involved in the regulation of gene expression [1–3], as well as genomic imprinting [4], X-chromosome inactivation [5], and the maintenance of genomic stability during mitosis and cell differentiation [6–8]. Variation in DNAm has been associated with many diseases, in particular cancers [9, 10], but also common disease [11] such as Parkinson’s disease [12], and rheumatoid arthritis [13]. Both genetic [14, 15] and environmental [16–18] factors are highly influential to the variation in DNAm levels across the genome. Studying the genetic architecture of DNAm can help us to understand the genetic control of DNAm and potential mechanisms through which genetic variants can affect complex traits via effects on DNAm.

Variation in DNAm levels is known to be under partial genetic control; a family based study estimated the average heritability of DNAm levels to be 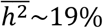 [15], whilst another study estimated the average SNP-based heritability to be 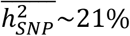 [19]. DNA methylation quantitative trait loci (mQTL) analyses have discovered many associations between common genetic variants and DNAm levels across the genome [14, 19-22]. Regional control of DNAm has been observed in regions of up to 3kb, through shared mQTL and correlations between DNAm levels across the region [14, 23], while a Bayesian co-localisation study found evidence for a shared genetic effect between ~282,000 pairs of CpG-sites at a median distance of ~110kb [22]. Overlap between mQTL and gene expression QTL (eQTL) has also been observed [14, 21], with genetic variants found to affect DNAm and gene expression levels pleiotropically [22, 24]. These observations point towards a possible mechanism through which genetic variants can alter gene expression levels via underlying differences in DNAm levels in a region.

Rare genetic variation has been shown to be important in the genetic architecture of complex traits, and gene expression [25–28]. Difficulties in studying the effect of rare variants reflect lack of power in traditional genome-wide association studies (GWAS) [29, 30]. Very large sample sizes are needed to detect statistically significant associations with rare variants given empirical estimated effect sizes. Various statistical methods have been developed to detect rare variant associations, gaining power by aggregating the effects of multiple rare variants, or looking for unusual variances in the effect sizes of rare variants in a region [31–37]. Using one of these rare variant association tests, there has been evidence of effects from rare variants on DNAm levels, even when there is no common variant association at the relevant CpG-site [38]. Rare variants have also been found to be enriched near the transcription start site (TSS) of genes in individuals with outlying levels of gene expression, particularly in those individuals with outlying levels of gene expression across multiple tissue types [27]. Other studies have found that the number of rare alleles within a region of the TSS of genes is on average higher in those individuals with lower, or higher levels of methylation than the population average, in both humans [39] and maize [40]. These rare variants are likely to be in promoter regions; hence, it is possible that they affect the DNAm levels in CpG-islands, which can have an effect on the gene expression levels [1, 41].

In this study, we investigate the effect of rare genetic variation on DNAm levels across the genome, and how DNAm levels may affect gene expression levels at nearby genes. We hypothesise that, similar to the association found between rare variants and outlying gene expression levels [27, 39, 40], there are associations between rare variants and outlying levels of DNAm. Outliers in DNAm have been associated with common diseases such as motor neurone disease [42] and type I diabetes [43], understanding the underlying mechanisms may help in determining the genetic etiology of these associations. In addition, CpG-sites are known to be highly mutable, with the mutation rate at CpG-sites estimated to be one order of magnitude higher than anywhere else in the genome, which results in an enrichment of mutations at CpG-sites in the genome [44, 45]. Knowing how mutations at CpG-sites will affect DNAm and gene expression levels in the genome may also be important for understanding the genetic etiology of complex trait diseases and cancers.

## Results

An overview of the methods used in this study, with the different data available to us is given in Figure 1.

**Figure 1.**
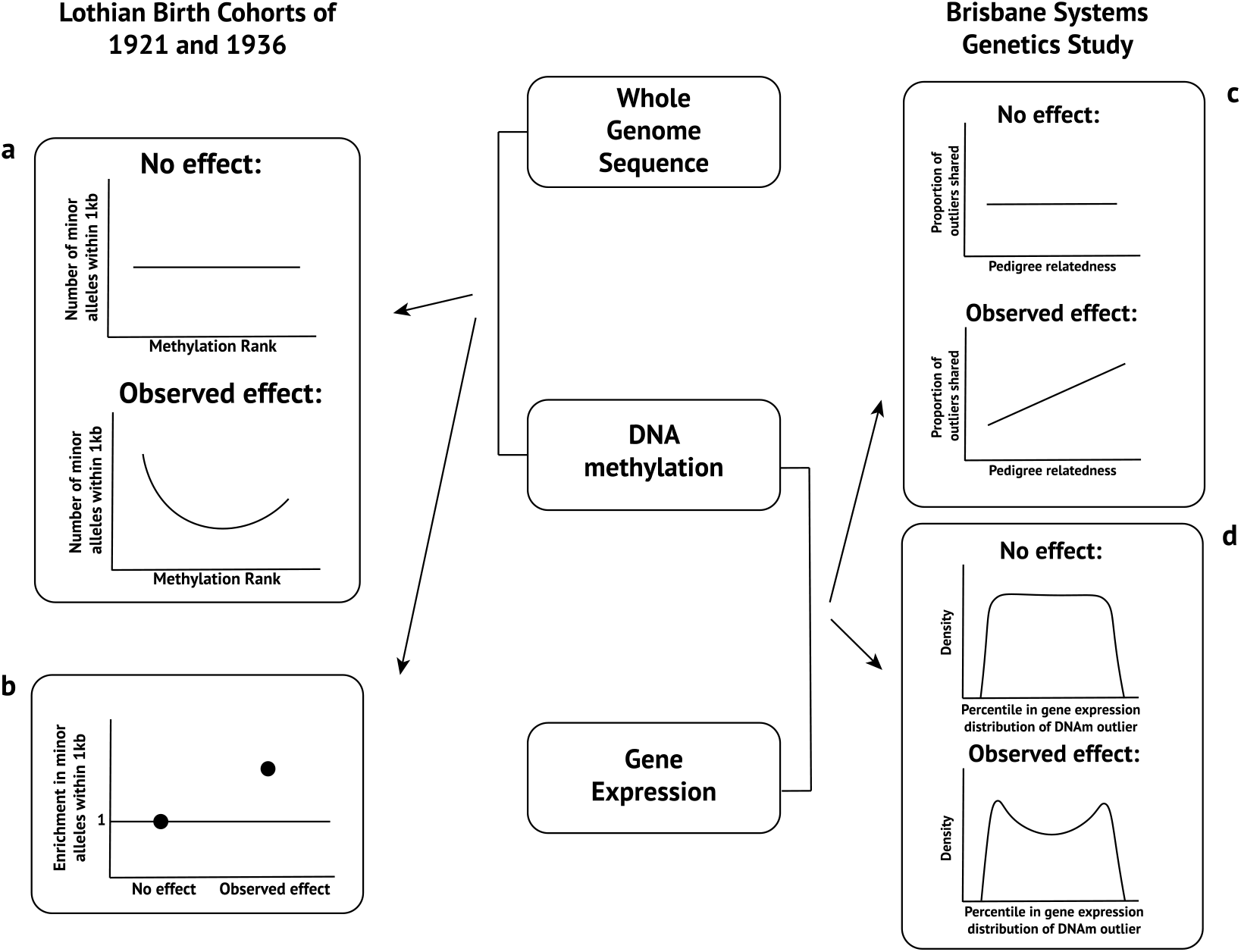
Overview of the methods used in this study. The Lothian Birth Cohorts of 1921 and 1936 were used to investigate the effect of genetic variants on DNA methylation levels, while the Brisbane Systems Genetics Study was used to examine the effect of DNA methylation levels on gene expression levels. In subfigure a, the number of minor alleles within 1kb is plotted against methylation rank (The individual with the n^th^ lowest DNAm levels will have a methylation rank of n at that CpG-site); in the case of no effect of genetic variants on DNAm levels, a uniform distribution is expected, any deviation from the uniform distribution is evidence for a genetic effect on DNAm levels. Subfigure b shows the enrichment of minor alleles within 1kb in outliers compared to non-outliers; in the case of no effect of genetic variants on DNAm outliers, the enrichment will be 1, any significant deviation from 1 is evidence of an effect of genetic variants on outliers of DNAm. In subfigure b, the proportion of outliers shared between pairs is plotted against the pedigree relatedness; if there is no genetic effect on DNAm outliers a slope of 0 is expected, any non-zero slope is evidence for a genetic effect on DNAm outliers. Finally, in subfigure d, the distribution of gene expression percentile of individuals with DNAm outliers at nearby probes is plotted; in the case of no effect from DNAm on gene expression, a uniform distribution is expected, any deviation from the uniform distribution is evidence for an effect of DNAm on gene expression.

### Detecting genome-wide genetic effects on DNA methylation

Using whole genome sequencing data and DNA methylation measures from the Illumina Infinium HumanMethylation450 array for n=1,261 individuals from the Lothian Birth Cohorts (LBC) of 1921 and 1936 [46], we tested for global effects of both rare and common genetic variants on DNAm levels across the genome. At each of the ~460,000 DNAm probes, individuals were ranked from lowest DNAm level to the highest, and the number of minor alleles within 1kb of the CpG-site were counted for each individual within a given minor allele frequency range. We then averaged the minor allele counts for each rank at each DNAm probe. If there is no genetic effect on DNA methylation for single nucleotide polymorphisms (SNPs) with a given allele frequency range, we would expect no relationship between the average minor allele count across ranks. We observe an inflation in allele counts at the lowest and highest ranks, for all MAF ranges (Figure 2), suggesting genetic effects from variants across all MAF ranges.

**Figure 2.**
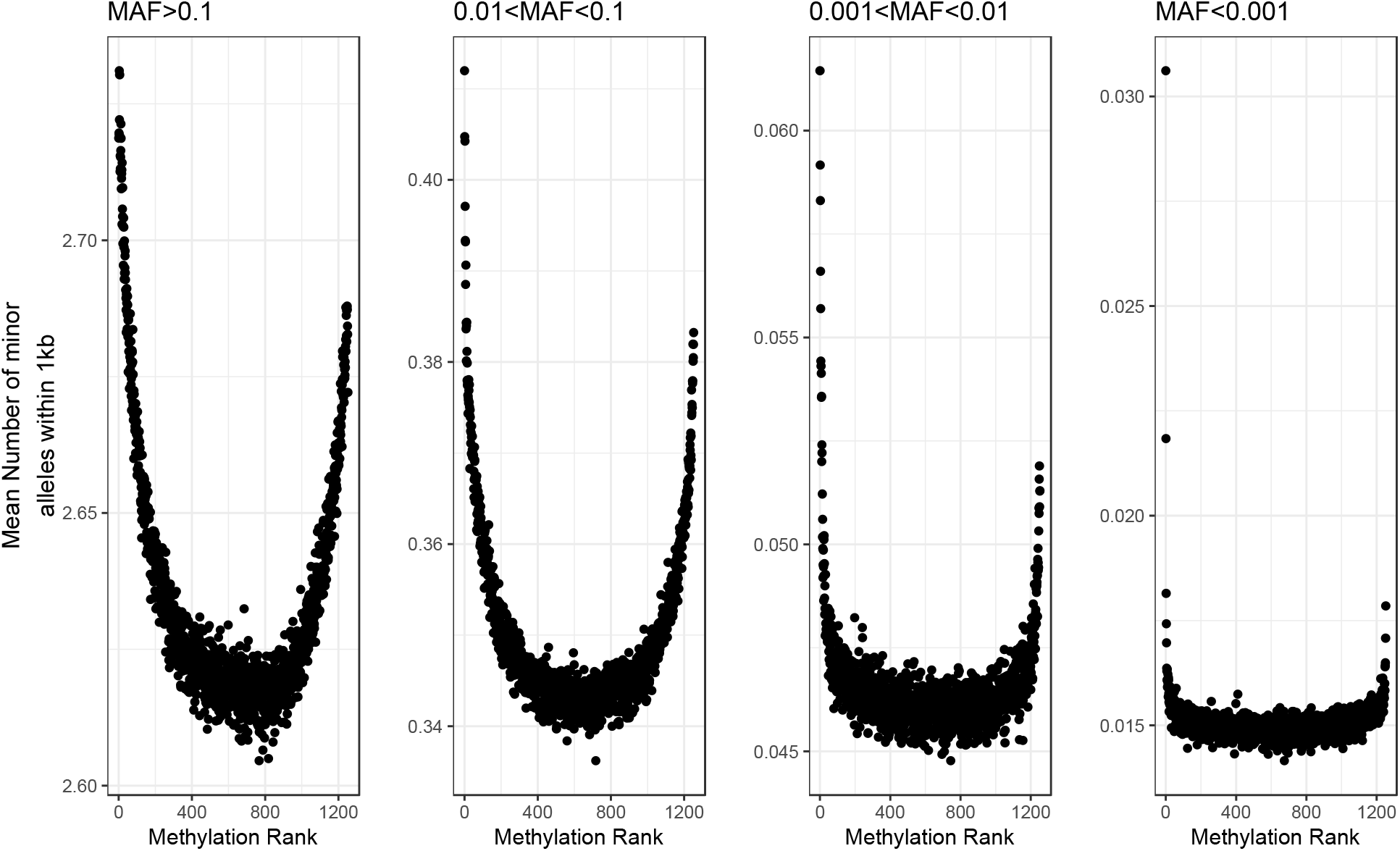
The mean number of minor alleles within 1kb of the CpG-site for each rank of DNAm levels across all autosomal probes. The analysis is split into 4 MAF bins. The inflation at the lowest and highest ranks is seen in each MAF bin, demonstrating that common and rare alleles both have an effect on DNAm genome-wide.

For the common variants (MAF > 0.1), we show that these effects are largely captured by mQTL analyses (Figure 3) by separating the ~50,000 probes with a significant mQTL detected in previous studies [20]. The inflation at the ends of the distribution remains for the DNAm probes with a known mQTL, while the majority of the inflation is removed for the remaining probes. This indicates that the majority of the relationship between methylation rank and SNPs for common variants is captured by known mQTL.

**Figure 3.**
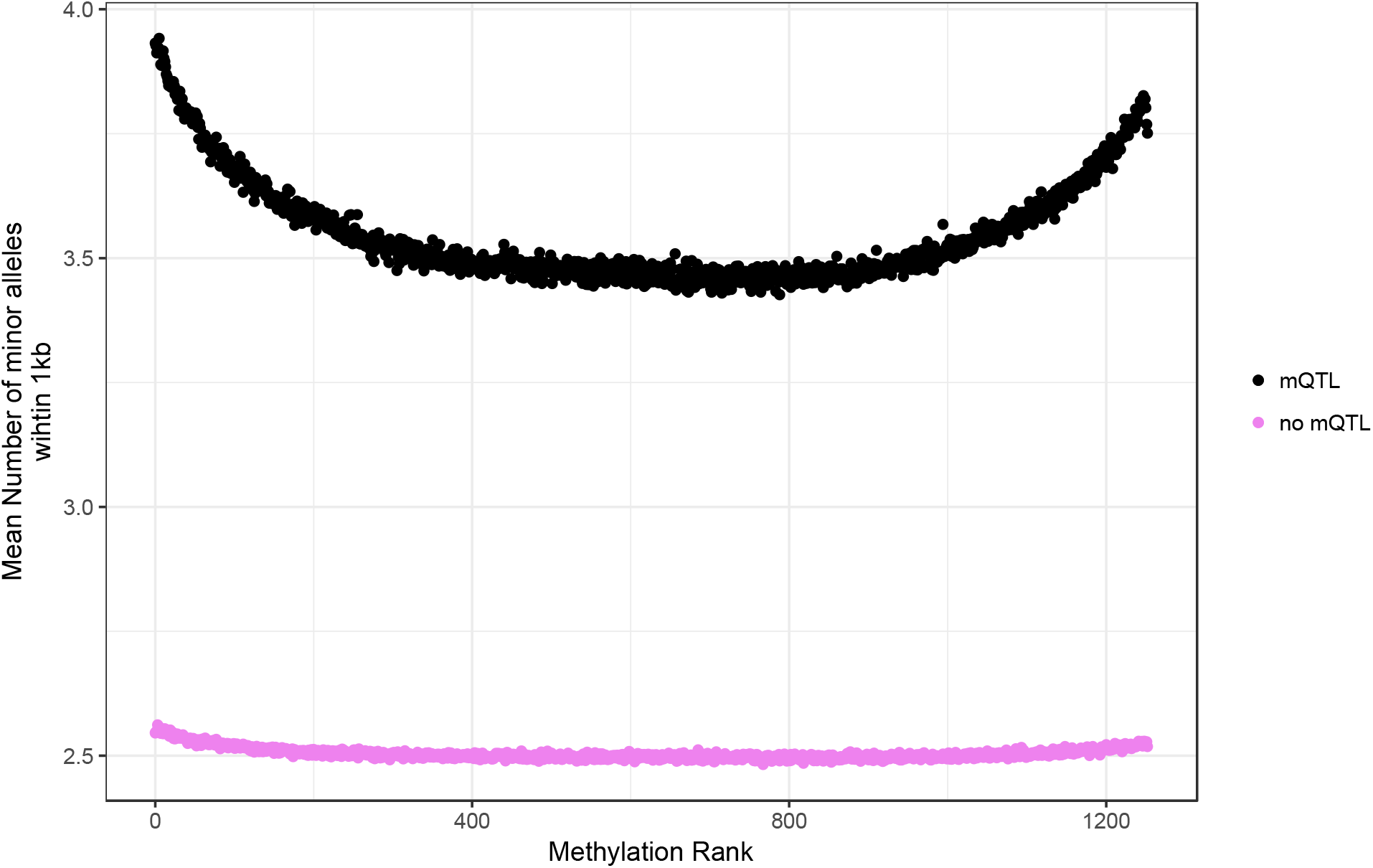
The effect of common genetic variants on DNAm is captured by mQTL analyses. Separating the ~50000 probes with a known mQTL from the remaining probes for the common variants (MAF>0.1), we see the inflation at the ends for the distribution is not as strong in the probes without an mQTL. There is also a mean difference of about 1 minor allele within 1kb, which is consistent with a nearby mQTL.

We also observe that the association between minor allele counts and methylation rank is not symmetrical, with the lowest ranks having a larger inflation than the highest ranks in the MAF bins. This observation suggests a bias towards SNP minor alleles decreasing DNAm levels across the genome. However, after separating the probes which contain a SNP at the CpG-site (CpG-SNP) from the rest of the probes, we see that the inflations are symmetrical for probes which do not contain a CpG-SNP (Figure 4). This suggests that the allele disrupting the CpG site is, on average, the minor allele, which may be attributed to a combination of bias in selection of CpG sites included on the array (sites which are generally CpGs were chosen), and a known mutational bias in the genome from (methylated) cytosine to thymine through the process of deamination [44]. We have shown that the effect of SNPs outside of the CpG-sites are approximately equally likely to increase or decrease DNAm levels (Figure 4).

**Figure 4.**
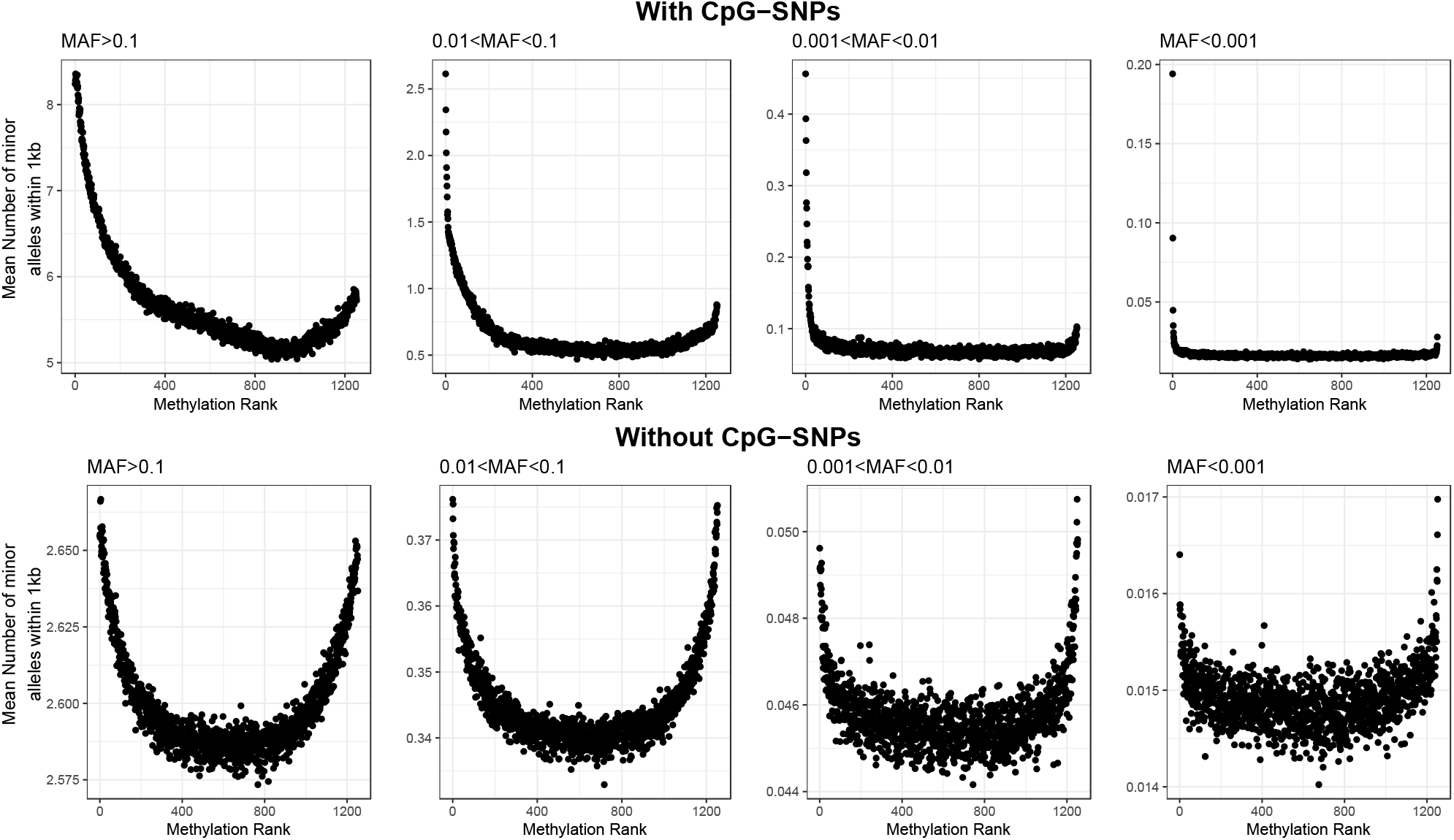
The mean number of minor alleles within 1kb of the CpG-site for each rank of DNAm levels across all autosomal probes with and without a CpG-SNP. The effects of CpG-SNPs were observed to reduce DNAm levels on average. On the other hand, the effects of SNP not at the CpG-site were observed to be symmetrical. This suggests that genetic effects outside the CpG-site are equally likely to increase or decrease DNAm levels.

While inflation in the minor allele count is observed for individuals with either lowly or highly ranked methylation values for all MAF classes, for the rare variants (MAF<0.001 and 0.001<MAF<0.01) we see that the inflation is largely restricted to the extremes of the distribution. This is consistent with rare variants driving more extreme levels of DNAm.

### Enrichment in rare alleles in individuals with outlying DNA methylation

We identified outlying DNAm levels at individual methylation probes using the subset of 642 individuals in the LBC dataset who have DNAm measurements at a minimum of three time-points. At a given time-point, an outlier was defined as a CpG-site in an individual with DNAm levels more than three times the interquartile range below the 1st quartile, or above the 3rd quartile at that CpG-site. We detected a total of 3,143,781 outliers in at least a single time-point of measurement (each individual can be outlying at multiple probes). Approximately 67% (309,114/459,309) of DNAm probes had at least one individual with outlying levels of DNAm. In addition, approximately 9% of the outliers at a CpG-site (281,311/3,143,781) were consistently outlying at that site across at least three time-points. The outlier burden (mean number of outliers per individual at a time-point [47]) was 2212 (out of 459,309 probes ~ 0.5%), reducing to 168 (~ 0.04%) when considering only those outliers stable across at least three time-points.

We observed an enrichment of ~1.2x the number of rare alleles (95% confidence intervals of [1.190, 1.222], [1.174, 1.193], and [1.157, 1.164] for variants with MAF<0.001, 0.001<MAF<0.01, and 0.01<MAF<0.1 respectively) within 1kb of the CpG-sites in individuals with outlying DNAm levels compared to individuals with non-outlying DNAm levels at all time-points (Figure 5). The enrichment in outliers remained statistically significant after removing the probes with a CpG-SNP (These probes may bias the enrichment as they will disrupt the methylation at the site which will likely result in outliers [48]). The enrichment of rare alleles in outliers compared to non-outliers stable across three to four time-points was larger ([1.356, 1.517], [1.363, 1.459], and [1.253, 1.288] in probes without a CpG-SNP, and [3.612, 3.994], [3.234, 3.4377], and [3.010, 3.083] in probes with a CpG-SNP for variants with MAF<0.001, 0.001<MAF<0.01, and 0.01<MAF<0.1 respectively) relative to the transient outliers observed to be outlying at a single time-point ([1.025 1.058], [1.028 1.047], and [1.030 1.038] in probes without a CpG-SNP, and [1.098 1.182], [1.116 1.166], and [1.134 1.155] in probes with a CpG-SNP for variants with MAF<0.001, 0.001<MAF<0.01, and 0.01<MAF<0.1 respectively. Figure 6).

**Figure 5.**
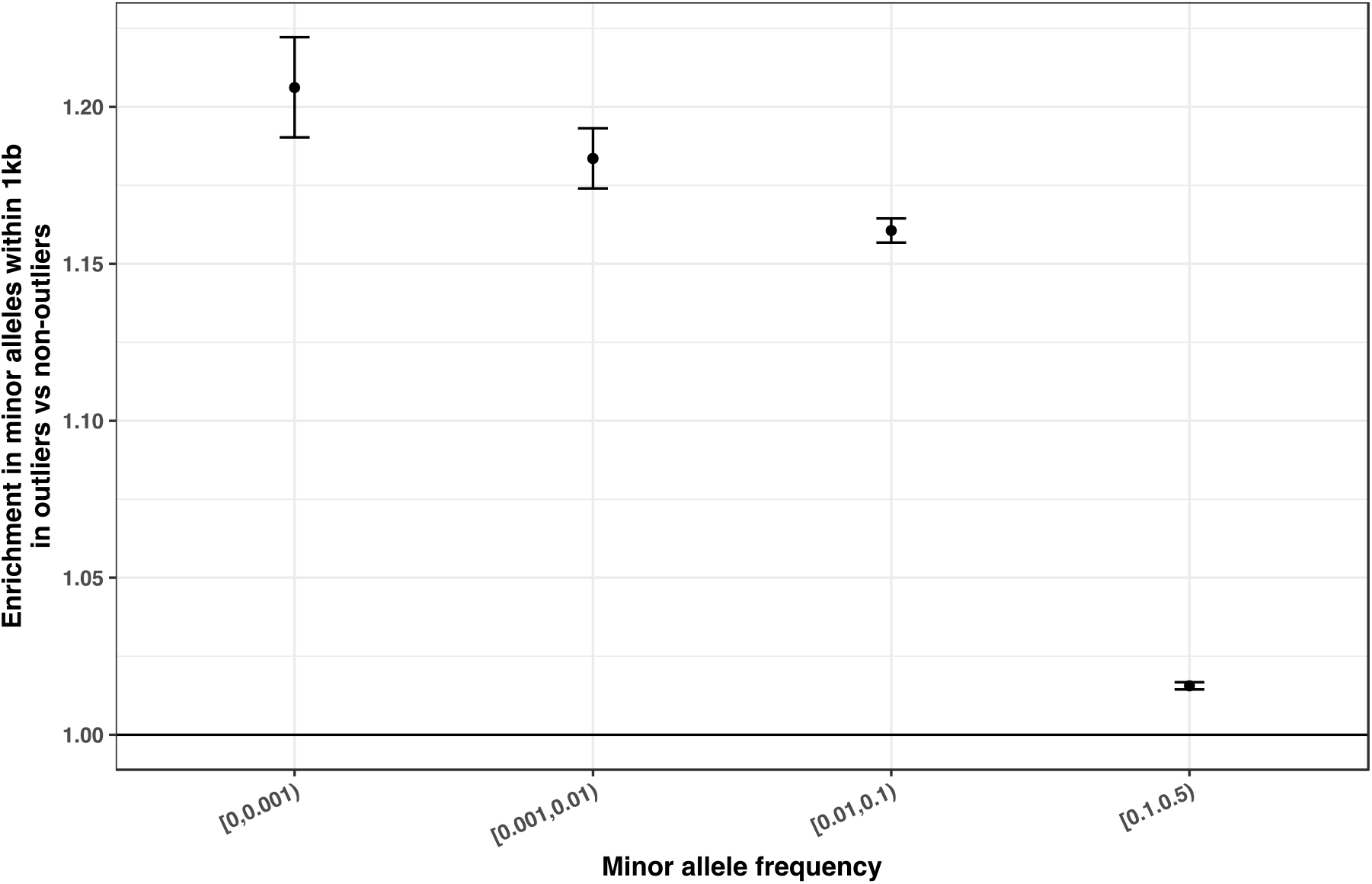
Outliers are enriched in rare alleles within 1kb of the CpG-site. The enrichment of minor alleles within 1kb of the CpG-site for individuals with outlying levels of DNAm levels compared to individuals with non-outlying levels of DNAm was significant for all minor allele frequency (MAF) groups.

**Figure 6.**
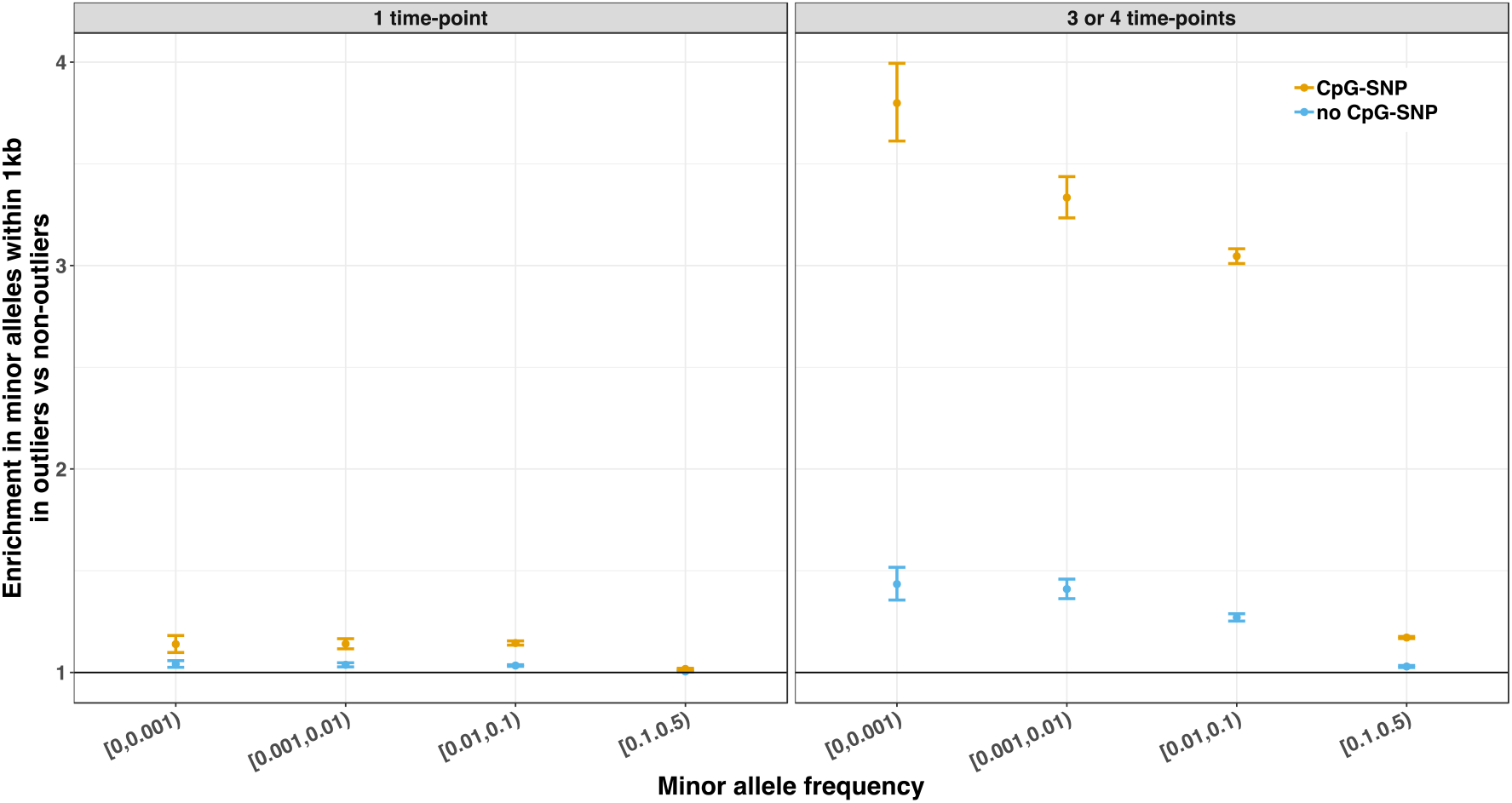
The enrichment of minor alleles in outliers compared to non-outliers at probes with and without a CpG-SNP. The enrichment in common alleles is not significant when excluding the probes with a CpG-SNP. For rare alleles, the enrichment in outliers remains significant in the individuals DNAm levels outlying stably across time, whereas the enrichment in individuals with DNAm levels outlying at only a single time-point is not significant.

### Outliers in gene-expression and DNA methylation are shared between relatives

Using the Brisbane Systems Genetics Study (BSGS) dataset [49] (n=595), which includes 67 MZ twin pairs, as well as many siblings and parent-offspring pairs with DNAm and gene expression array data, we detected a total of 1,481,297 outliers in DNAm levels (using the same definition of outliers as before), and 446,916 outliers in gene expression levels (using the definition of outliers as a gene expression probe in an individual with gene expression levels outside of 1.5x the interquartile range of the 1^st^ or 3^rd^ quartile).

We observed a linear relationship between the proportion of DNAm outliers (R^2^=0.52, slope=0.31, and p<10^−323^) and gene expression outliers (Adjusted R^2^=0.02, slope=0.03, p<10^−323^) shared between each pair of individuals, and their pedigree relatedness (Figure 7). This is consistent with genetic effects underlying outlying levels of DNAm levels, as well as gene expression levelss across the genome. However, there was very little overlap between gene expression outliers and DNAm outliers, with 6.1% of individuals with a gene expression outlier also having a DNAm outlier at the nearest annotated gene.

**Figure 7.**
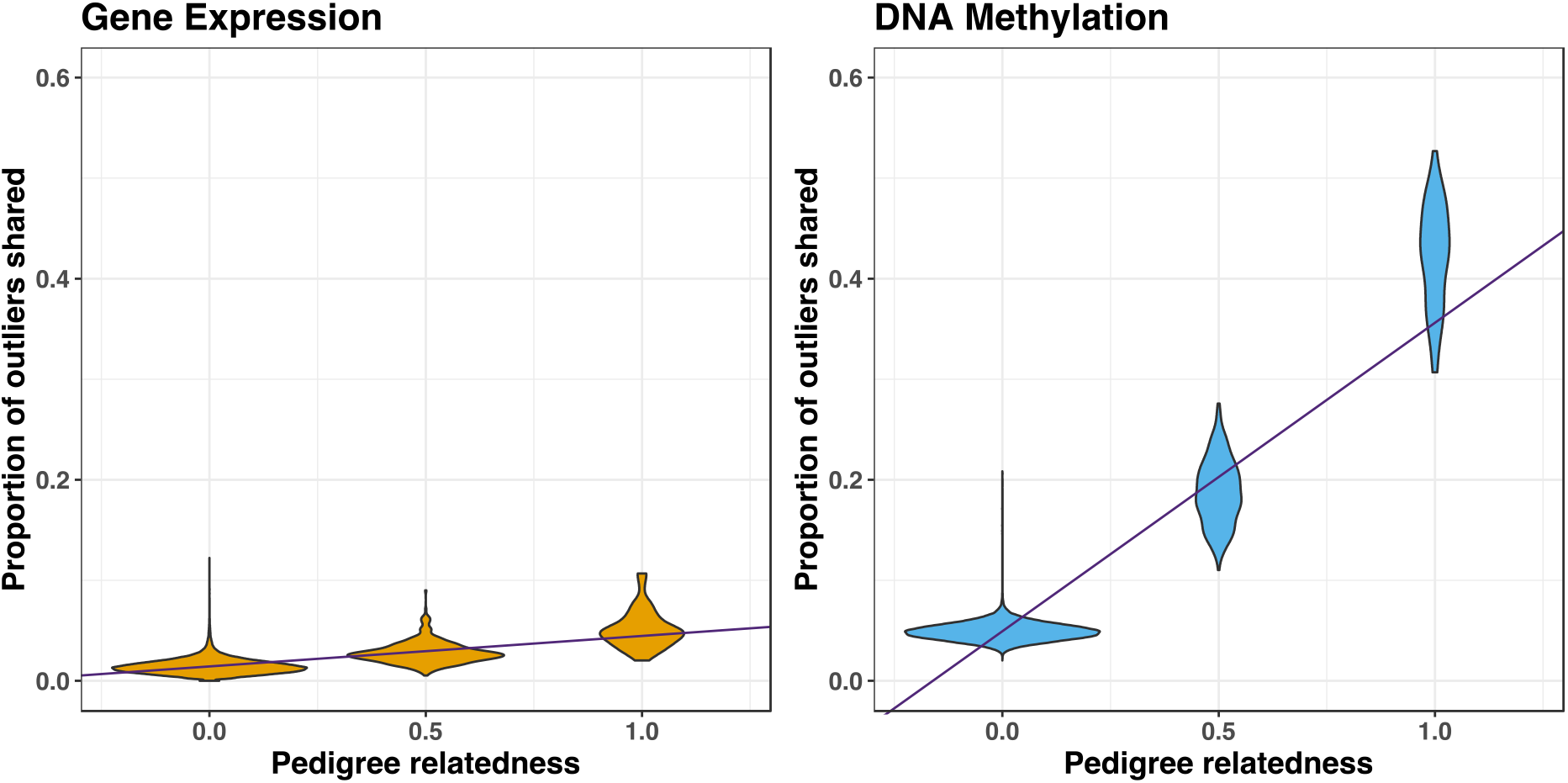
Outliers in DNAm, and gene expression are shared between relatives more often than at random. The linear relationship between pedigree relatedness and proportion of outliers shared suggests a genetic component to the outlying levels of DNAm and gene expression. The difference in slope suggests a stronger genetic effect on the DNAm levels compared to gene expression levels.

### Outlying levels of DNA methylation are associated with a change in gene-expression

Although the overlap of outlying DNAm and gene expression was not substantial, we tested whether the outlying DNAm levels correlates with any change in gene expression levels. For individuals with outlying levels of DNAm at a CpG-site, if the DNAm levels have no effect on gene expression levels, we would expect those individuals to be uniformly distributed across the gene expression distributions. Firstly, we paired DNAm probes to gene expression probes using significant common variant co-localisation established using a summary data-based Mendelian randomisation (SMR) study [24]. The rank of gene expression levels for individuals with outlying methylation levels at SMR-linked probes showed significant deviance from the uniform distribution (Kolmogorov-Smirnov one sample test D=0.03, p<10^−323^, Figure 8), indicating an association between outlying levels of DNAm levels on gene expression levels.

**Figure 8.**
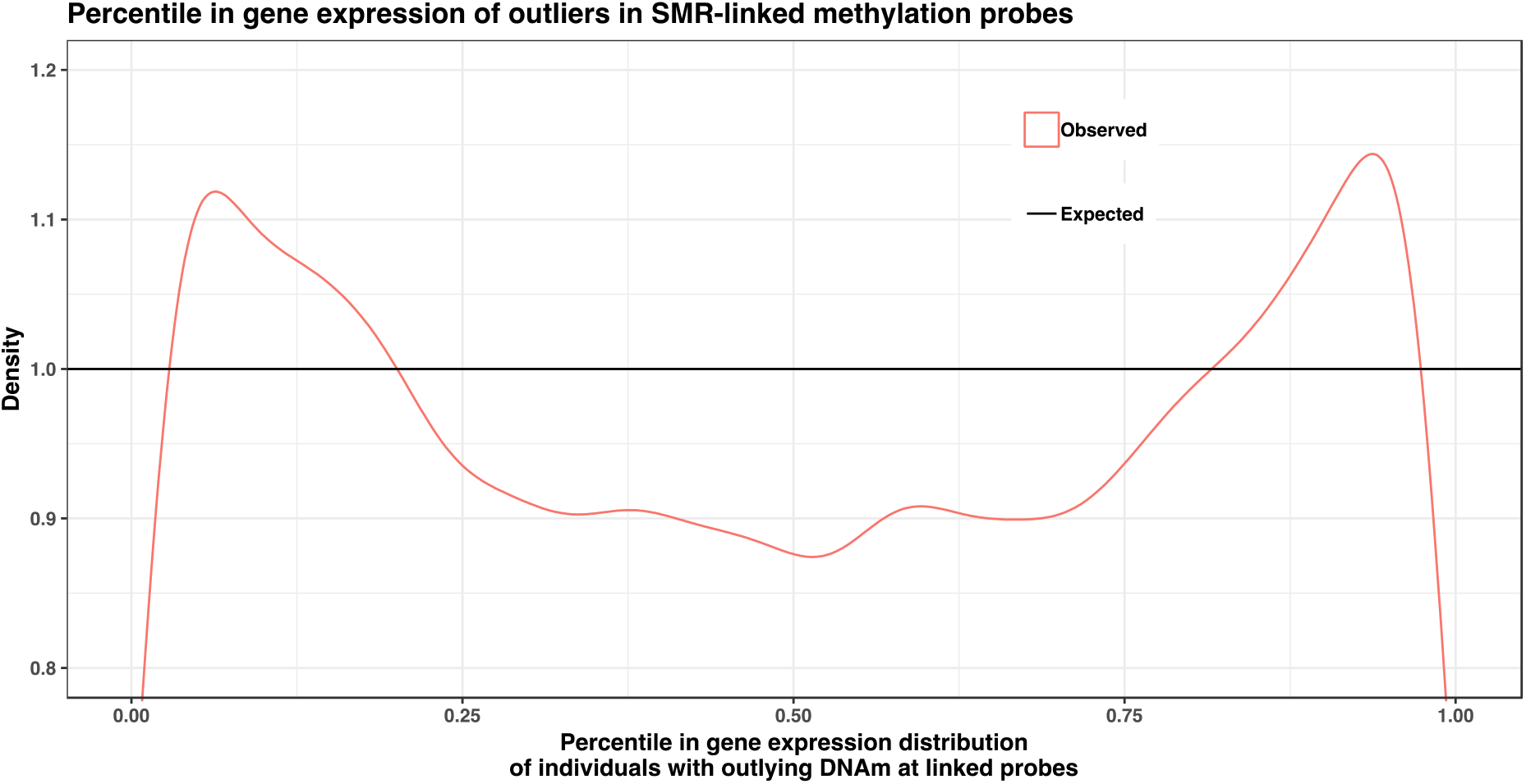
Density plot of the percentile in the gene expression distribution of individuals with outlying DNAm levels at a linked DNAm probe. Taking all DNAm and gene expression probe pairs linked through a summary-data based Mendelian randomisation analysis, we observe a significant deviation from the uniform distribution (Kolmogorov-Smirnov one sample test D=0.03 and p<10^−323^), suggesting that outlying levels of DNAm are associated with a change in the gene expression levels.

Secondly, we relaxed the criteria for linked DNAm and gene expression probes, using a distance-based pairing, taking all probe pairs within 10kb of each other. This introduced more noise into the analysis as not all DNAm and gene expression probes will be linked in any way. However, we still observed a significant deviation from the uniform distribution (Kolmogorov-Smirnov one sample test D=0.006, p<10^−323^, Figure 9). These results correspond to a correlation between outlying levels of DNAm and a change in gene expression levels at the relevant genes.

**Figure 9.**
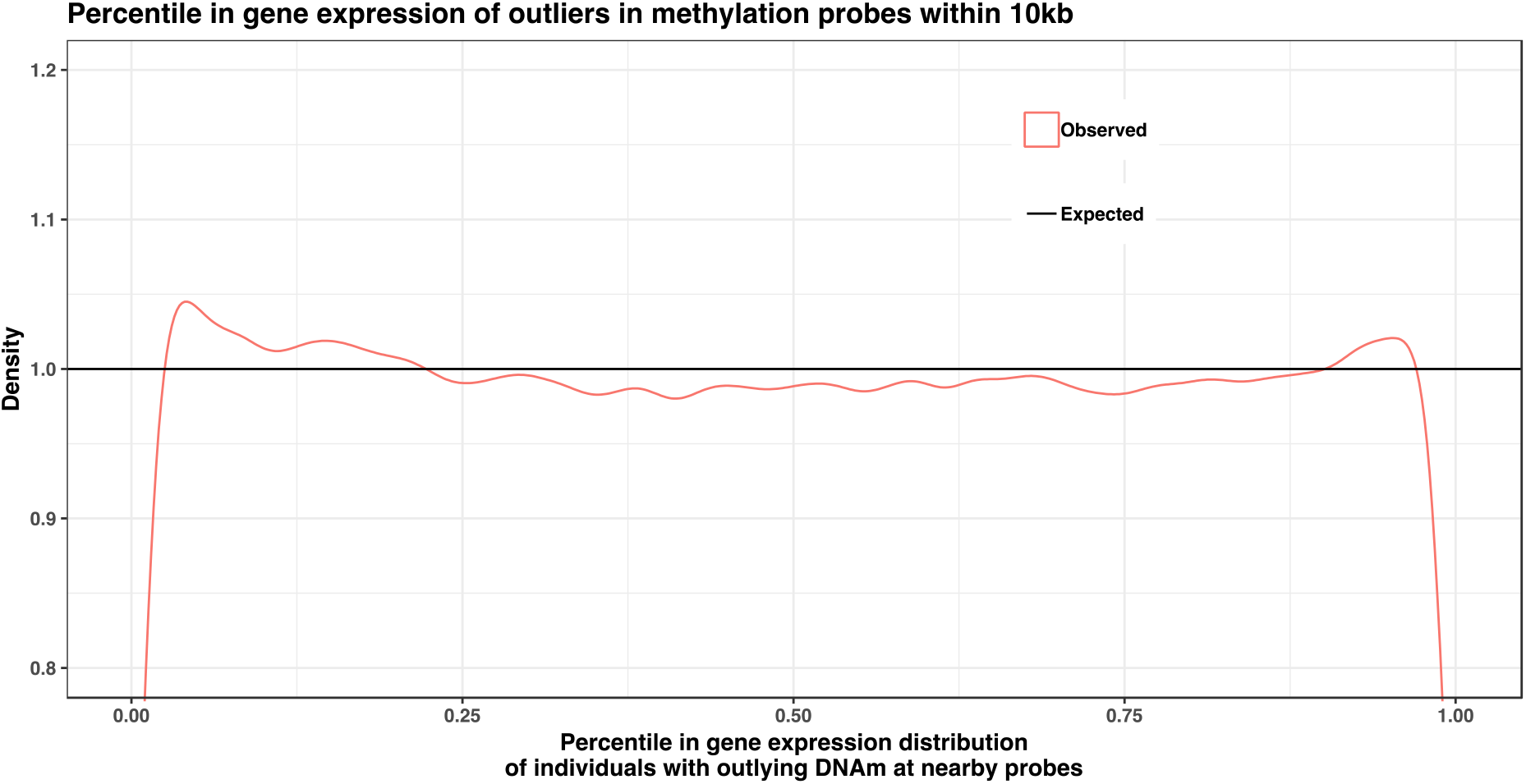
Density plot of the percentile in the gene expression distribution of individuals with outlying DNAm levels at a DNAm probe within 10kb. Taking all individuals with outlying DNAm levels at DNAm probes within 10kb of a gene expression probe, we observe which percentile they lie in the gene expression distribution at the gene expression probe. We observe a significant deviation from the uniform distribution (Kolmogorov-Smirnov one sample test D=0.006 and p<10^−323^), suggesting that outlying levels of DNAm are associated with a change in the gene expression levels.

## Discussion

This study examined the links between DNAm levels, rare genetic variants, and gene expression levels across the genome. We combined multiple lines of evidence to demonstrate the role of rare variants in outlying DNAm levels. Outlying levels of DNAm are further demonstrated to be associated with gene expression levels at nearby genes.

We examined the patterns of effects from common and rare genetic variants, within 1kb of the CpG-site, on DNAm levels across the genome. We found that rare alleles were associated with extreme levels of DNAm. In addition, we observed a significant enrichment of rare alleles within 1kb of CpG-sites in individuals with outlying levels of DNAm compared to individuals with normal DNAm levels at that CpG-site. Our results suggest that, in addition to common variants, rare variants also play a role in the control of DNAm levels across the genome.

DNAm levels at many CpG-sites are known to be correlated with age [23, 50], and changes in environment are also known to have an effect across time [16–18]. In our analysis, we found that outliers in DNAm levels which are present at only one time-point had almost no enrichment for rare alleles within 1kb of the CpG-site compared to non-outliers, but those probes outlying across multiple time-points within an individual had significant enrichment. This result suggests that transient outliers detected at a single time-point (2586888/3134194 ≈ 83% of the outliers in our study) are likely caused by environmental effects or measurement error, but the outliers stable across time are more likely to have an underlying genetic cause. This genetic effect underlying outliers in DNAm was confirmed using a family study design in an independent dataset. This is consistent with previous observations made using the LBC dataset in Shah et al. [51] who noted that many CpG-sites across the genome had stable DNA methylation across the lifetime, and these results are also in concordance with the observation made by Gaunt et al. [19] that the majority of mQTL are stable across time.

Similar to aggregation tests, we looked at enrichments and not associations with individual variants (which would be difficult to detect due to the power needed to reach statistical significance) we cannot say which variants have an effect and which do not. Notwithstanding, only a single rare variant (MAF<0.01) was observed within 1kb of the CpG-site in over 19% (25,591/131,903) of the outliers that were stable across time and had no CpG-SNP. However, even in these cases of only one rare allele within 1kb, we cannot determine causality without functional experiments.

Previous studies have found correlations between DNAm and gene expression, and an overlap in the association of common genetic variants between them [14, 21, 41, 52-55]. In this study, we show that outliers in DNAm levels are associated with a change in gene expression levels at nearby genes. Summary-data based Mendelian randomisation [56] analyses have provided us with evidence of pleiotropic effects of common variants on DNAm and gene expression levels across the genome [22, 24]. In addition, the proportion of phenotypic variance explained by the lead variant at a mQTL was, on average, larger than the phenotypic variance explained by the same variant at a co-localised eQTL and at a co-localised higher-order complex trait QTL, such as height [24]. This attenuation in effect size of the variant at each step suggests a mechanism of effect from genetic variant to DNAm, to gene expression, to higher-order complex trait. In this study, we observed that large differences in DNAm often corresponded to smaller differences in gene expression, which would fit into this hypothesised directional mechanism of effect. In addition, the difference in slope in Figure 7 also suggests a larger effect from genetic variation on DNAm levels, than gene expression levels. This mechanism may be important to consider, as DNAm has been shown to be associated with many common diseases [11], and as methylation outliers are relatively easy to detect, it could provide a useful tool for future research.

A limitation of our study was that of the two data sets available to us, one (LBC) had WGS and DNAm array data, whereas the other (BSGS) had SNP array, DNAm array and gene expression array data. Ideally the study would be conducted on a cohort with all data types. With the increasing availability of whole genome sequence data, as well as RNA-seq and DNAm array/bisulfite sequence data, a more comprehensive study of the effects of rare variants on both DNAm and gene expression would provide a better understanding of the mechanisms underlying genetic effects on complex traits. Other epigenetic mechanisms, such as histone tail modifications, are highly correlated with DNAm levels, are under shared genetic control [14, 21], and are also involved in the regulation of gene expression [54, 57]. We hypothesise that other epigenetic modifications may also show similar patterns of effects to what we found in DNAm, and including these into future analyses could potentially provide a more complete picture of the shared genetic control between DNAm, other epigenetic modifications, and gene expression.

In summary, this study provides a novel insight into the effect of rare variants on DNAm levels across the genome, and shows that extreme differences in DNAm are associated with gene expression levels at nearby genes, which may be driven by rare genetic variation.

## Methods

### Lothian Birth Cohorts of 1921 and 1936

The Lothian Birth Cohorts of 1921 and 1936 (LBC) [46] are part of a longitudinal study of cognitive ageing. DNA was extracted from whole blood samples from which DNAm levels were measured using the Illumina HumanMethylation450 BeadChip array across three or four time-points. The raw intensity data were background corrected, corrected for cell-type and quantile normalised using standard QC protocols, and the DNAm beta-values were generated using the R package *meffil* [58].

DNAm levels were measured at an average age (sd) of 79.1 (0.6), 86.7 (0.4), and 90.2 (0.1) years in the LBC1921 cohort and ages 69.6 (0.8), 72.5 (0.7), 76.3 (0.7), and 79.3 (0.6) years in the LBC1936. Of the 1342 individuals with DNAm measured at one point, 642 had at least three timepoint measurements. While DNAm levels across the genome are known to change with age [23, 50], this is not a confounding factor in our analysis as the age ranges within each wave of measurement are very narrow (mean standard deviation of age for each cohort in each wave was 0.6 years).

Whole genome sequencing was performed on the HiSeq X with an average coverage of 36x (minimum 19.6x, maximum 65.9x). Details of the QC can be found in Prendergast et al. 2019 [59]. Briefly, reads were mapped using BWA [60] to the build 38 of the reference genome, and GATK [61] was used for variant calling. Variant effect predictor (VEP) [62] was used to annotate variants and gene models from the version 85 release of Ensembl.

### Brisbane Systems Genetics Study

The Brisbane Systems Genetics study (BSGS) [49] was a dataset designed to study the genetic effects on gene expression, and the role of gene regulation in complex traits. DNAm levels were measured, in whole blood using the Illumina Infinium HumanMethylation450 BeadChip array, on 614 individuals from 117 families, including monozygotic twin pairs, dizygotic twin pairs, sibling pairs, and parents. The QC of the DNAm data was performed using the same pipeline as with the LBC data. gene expression levels were measured in whole blood on 846 individuals using the Illumina HumanHT-12 v4.0 BeadChip array. The QC of the gene expression data are detailed in Lloyd-Jones et al. 2017 [63]. Briefly, the gene expression levels were normalised using variance stabilization [64], quantile normalised using the *limma* software [65], followed by PEER factor adjustment [66], with 50 factors, correcting for covariates such as age, sex, cell counts, and batch effects. Both DNAm and gene expression levels were measured on a total of 595 individuals.

An overview of the methods used to investigate the effects of genetic variants on DNAm levels and gene expression levels using the LBC and BSGS datasets are shown in Figure 1.

### Detecting genome-wide effects on DNAm

Following similar procedures to Zhao et al. [39], and Kremling et al. [40], we ranked the individuals in the LBC data at each DNAm probe from lowest DNAm beta-value to the highest, and counted the number of minor alleles within 1kb of the CpG-site for each individual. We averaged this value at each rank across all autosomal probes to get the mean number of minor alleles within 1kb of a CpG-site. We did this for 4 MAF ranges, MAF>0.1, 0.1>MAF>0.01, 0.01>MAF>0.001, and 0.001>MAF, which allowed us to separate the effects of common and rare variants. The rarest MAF bin (MAF<0.001) corresponded to variants with one or two observed minor alleles in our dataset. This analysis was performed using the 1^st^ wave of measurements in the LBC dataset to maximise sample size.

### Detecting outliers

We defined DNAm outliers as a CpG-site in an individual with DNAm levels outside 3 interquartile ranges (IQRs) from the 1^st^ quartile (Q1) or the 3^rd^ quartile (Q3) of the DNAm levels at that CpG-site. The standard 1.5 IQRs from Q1 or Q3 compares to 3 standard deviation from the mean in a perfectly normal distribution. Our definition is slightly more stringent than this, as the distribution of DNAm levels can be highly skewed. For detecting outliers in the gene expression data, which had more symmetric distributions, the standard 1.5 IQR from Q1 and Q3 definition was used.

### Enrichment of rare alleles around CpG-sites

We defined enrichment as,

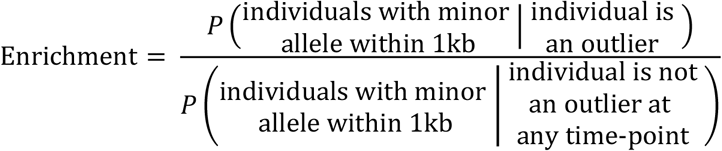

In words, we defined enrichment as the probability of an individual having a minor allele within 1kb of a CpG-site given they have outlying DNAm levels at that site, divided by the probability of an individual having a minor allele within 1kb of a CpG-site given they don’t have outlying DNAm levels at that site. This is similar to the definition used in Li et al. [27], although they used a slightly different definition of outliers (>2 standard deviations from the mean).

### Proportion of outliers shared

To compute the proportion of outliers shared between each pair of individuals, we used the formula 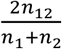, where n_1_ is the number of outliers for individual one, n_2_ is the number of outliers for individual two, and n12 is the number of outliers shared between the individuals. The relatedness coefficients were obtained from pedigree data.

### Testing for association between outlying levels of DNAm and gene expression

To test for an association between outlying levels of DNAm and gene expression, the percentile in the gene expression levels distribution at a gene expression probe was calculated for each individual with outlying DNAm levels at the paired DNAm probe. We used two methods to pair DNAm probes to gene expression probes. Firstly, we linked DNAm probes through a shared common variant co-localisation with the gene expression probe detected using the Summary-data based Mendelian Randomisation (SMR) method [24, 56]. We also used all pairings of gene expression probes within 10kb of the CpG-sites. This represents a trade-off between number of pairs included in the analysis and including pairs of gene expression and DNAm probes that have no biological connection beyond proximity. Under the null hypothesis of no association between outlying DNAm and gene expression levels, the rank of gene expression levels for individuals with outlying DNAm levels should be uniformly distributed. We tested for deviation from the uniform distribution using the Kolmogorov-Smirnov one sample test [67], which tests the degree of agreement between the sampled values and a theoretical distribution, in our case the uniform distribution.

